# Predicting current and future distribution of the common waxbill using a mechanistic modelling approach

**DOI:** 10.1101/2025.11.16.688699

**Authors:** Marina Sentís, Cesare Pacioni, Luís Reino, Luc Lens, Diederik Strubbe

## Abstract

Biological invasions and climate change are two of the most pressing drivers of biodiversity loss worldwide. Anticipating where invasive species are likely to establish, as well as the potential impact of climate change on their range expansion, is essential for early detection and targeted management. In this study, we use a mechanistic species distribution model (SDM) to evaluate the current and future areas at risk of invasion by the common waxbill (*Estrilda astrild*), a widespread avian invader. Our model accurately predicts the species’ current range in Iberia and identifies additional climatically suitable areas, particularly in southern and western Europe. Under warming scenarios of +2 °C and +4 °C, suitable areas expand northwards, with over two-thirds of Europe classified as suitable under the most extreme scenario. These results suggest that the species is already operating near the cold limits of its thermal niche in parts of its invasive range, and that rising temperatures may remove these constraints, allowing expansion into previously unsuitable areas. Our findings demonstrate the power of mechanistic models in identifying regions at risk of colonisation and underscore the importance of early intervention and targeted monitoring in areas projected to become suitable.

## 1. Introduction

Climate change and biological invasions are among the greatest threats to global biodiversity and ecosystem functioning (Bellard et al. 2012, 2016, IPBES 2023). They often interact synergistically, and as global temperatures rise and precipitation patterns shift, regions previously unsuitable for certain invaders may become climatically favourable, facilitating range expansion and increasing the likelihood of successful establishment (Hellmann et al. 2008, Mainka and Howard 2010, Shabani et al. 2020). At the same time, climate change can reduce habitat suitability and erode the adaptive capacity of native species, further tipping the balance in favour of invaders (Mainka and Howard 2010). Shifts in biotic and abiotic environmental conditions caused by climate change may not only facilitate more biological invasions, but also increase the difficulty of eradicating or managing established invasive populations (Hellmann et al. 2008, Colberg et al. 2024). Consequently, because removal of established invaders is costly (Leung et al. 2002, Diagne et al. 2020), early prevention measures are crucial, making predictions of where invasive species will occur essential for proactive conservation and management strategies (Bradshaw et al. 2016).

Mechanistic species distribution models are increasingly recognised as powerful tools for forecasting how species respond to novel environments and changing climates (Estrada et al. 2016, Kearney et al. 2021). By explicitly characterising a species’ fundamental niche, the full range of environmental conditions that are physiologically tolerable, these models offer a process-based approach to projecting species distributions based on biophysical principles. Unlike correlative SDMs, which infer environmental suitability from observed occurrence data, mechanistic models link organismal traits to environmental constraints, allowing for predictions grounded in physiology rather than statistical associations (Kearney et al. 2009, Porter and Kearney 2009). Correlative SDMs have been widely used and have proven valuable in many contexts (Guisan et al. 2013, Srivastava et al. 2019), particularly for estimating the realised niche based on current distributions (Araújo and Peterson 2012, Petitpierre et al. 2017). However, their assumption that species are in equilibrium with their environment (Thuiller et al. 2005) may be challenged in the case of biological invasions or under rapid climate change. Introduced species can undergo niche shifts in novel environments (Hill et al. 2017, Atwater et al. 2018, Liu et al. 2020, Bates and Bertelsmeier 2021), and climate change may generate environmental conditions that lie outside the range of current observations. In such cases, the predictive accuracy of correlative SDMs may be compromised (Buckley et al. 2010, Sinclair et al. 2010).

Because mechanistic models estimate the fundamental rather than the realised niche, they are often argued to be less affected by niche shifts, and they can be applied even when occurrence data are sparse or absent, as is often the case for emerging invasive species (Urban et al. 2016, Bates and Bertelsmeier 2021). This should make them particularly suited to invasion risk assessments and forward-looking applications under climate change. While a common limitation of mechanistic models has been their requirement for detailed, species-specific trait data (Jeschke and Strayer 2008, Buckley et al. 2010), this barrier is gradually being lowered by the expansion of global trait databases and the development of standardised trait protocols (Bennett et al. 2018, Herberstein et al. 2022, Tobias et al. 2022).

To test whether a refined biophysical (mechanistic) approach, NicheMapR, can address limitations of previously published correlative and mechanistic SDMs, here we focus on the common waxbill (*Estrilda astrild*). Common waxbills are small estrildid finches native to sub-Saharan Africa that have become globally widespread, primarily via the pet trade (Carrete and Tella 2008, Stiels et al. 2011). They belong to the Estrildidae family, which includes some of the most frequently traded and successful avian invaders (Su et al. 2022). In Europe, the common waxbill was first recorded as established in Portugal in the 1960s and has since expanded its range across the Iberian Peninsula, particularly in irrigated agricultural landscapes (Reino and Silva 1998, Cardoso and Reino 2018). It is a generalist granivore with flexible foraging behaviour, often forming large flocks in open habitats (Ribeiro et al. 2020). Despite its widespread establishment, no significant negative impacts on native biodiversity have been documented (Reino et al. 2009). Several studies have already developed correlative species distribution models (SDMs) to estimate waxbill habitat suitability across both native and invasive ranges. However, model performance has been shown to be sensitive to the choice of input data and modelling approach (Silva et al. 2002, Reino et al. 2009, Sullivan et al. 2012, Strubbe et al. 2013, Sullivan and Franco 2018, Baquero et al. 2021). More recently, Santana et al. (2024) employed a high-resolution temporal dataset of waxbill occurrences in the Iberian Peninsula to evaluate SDM performance across different invasion stages. They found that models trained with early-stage invasion data had limited predictive power. However, iteratively updating SDMs as the invasion progressed and more occurrence data became available led to improved model accuracy in predicting the species’ potential invasive range. Despite this improvement, such a reactive modelling approach is limited in its capacity to support proactive management, as it depends on the availability of new data following range expansion.

Strubbe et al. (2023) applied a mechanistic SDM to the common waxbill in Europe, but relied on ecophysiological and morphological trait values derived from the literature and a small number of museum specimens, and used standard, coarse-grid bioclimatic averages to characterize the microclimates used by waxbills, resulting in comparatively poor predictive performance for the common waxbill. These shortcomings may have compromised the model’s ability to capture the common waxbill’s physiological tolerances in different environments. To address these, here we develop a detailed biophysical model that incorporates empirical trait measurements (e.g., metabolic rates and feather characteristics) to more accurately capture the fundamental niche of the common waxbill, using high-resolution estimates of climate and weather variables for microclimate modelling. Our study leverages recent advances in species-specific ecophysiological data, including measurements of metabolic rate and body temperature, alongside detailed morphological trait data (e.g., body size and plumage characteristics) collected from a large number of museum specimens and wild individuals (Pacioni et al. 2023, Sentís et al. 2023, 2025). Forecasts of invasion risk are then made using a previously parameterized and validated NicheMapR model, which has been shown to accurately capture the energy expenditure of captive waxbills (Sentís et al. 2023). We first assess the invasion risk of waxbills across Europe under current climate conditions, using occurrence data from their invasive range to evaluate the accuracy of our mechanistic model. We then forecast how invasion risks may shift under two warming scenarios (+2 °C and +4 °C).

## 2. Material and methods

### Microclimate model

As it is the microclimate—not the broader macroclimate—that governs the thermal and moisture conditions encountered by organisms (Kemppinen et al. 2024), we employed the microclimate model from the ‘NicheMapR’ modelling platform (Kearney and Porter 2017) to simulate microclimates experienced by waxbills, using the R software v. 4.3.2 (R Core Team 2023). Specifically, we used the ‘micro_terra’ function to integrate the high-resolution (∼4 km) TerraClimate dataset, which provides monthly climate data from 1958 to 2020, along with future projections representing +2 °C and +4 °C of global warming (Abatzoglou et al., 2018). NicheMapR generates hourly estimates of key environmental parameters at the average height of the modelled animal, including air temperature, wind speed and relative humidity. Since common waxbills are predominantly granivorous and forage close to the ground in search of grass seeds, we modelled microclimates at a height of 0.9 m. Microclimate conditions were modelled for 12 days at a 5 x 5 km resolution across Europe, representing mid-month conditions for each calendar month. Annual microclimate outputs were generated for each year between 2010 and 2020, and the resulting suitability rasters were averaged across this period to obtain a single mean suitability raster.

Seasonal surface albedo values were derived from the Landsat 8 Surface Reflectance data. Cloud-free observations for January-December 2015 were selected, and reflectance values were extracted. To calculate albedo, we used the equations given by (Liang 2001, Smith et al. 2023). Windspeed and roughness height (RUF) values were derived from a land cover dataset at 30 m resolution (Parente et al. 2021). Different reduction percentages were applied to windspeed depending on vegetation types and landscape features. For example, reductions of 80% were assigned to broad-leaved forests (Davies-Colley et al. 2000) and 75% for wetlands (Warrilow et al. 1978). Roughness height modifiers, as well as maximum shading percentages, were similarly adjusted according to the land cover categories (Silva et al. 2007). These parameters were incorporated dynamically for each pixel to reflect spatial variability in windspeeds and shade. A digital elevation model (DEM) was obtained from WorldClim’s global elevation dataset (Fick and Hijmans 2017). Slope and aspect were computed from the DEM using the terrain() function from the ‘terra’ package. These were then integrated into the microclimate model to account for terrain-specific variations. All other microclimate model parameters were kept at their default values. The model outputs provided hourly estimates of air temperature at reference height and at the height of the birds (°C), sky and surface temperatures (°C), wind speed (m s^−1^), relative humidity (%), and solar radiation (W m^-2^). These data were incorporated into the biophysical model described below.

### Ecophysiological model

We used the endotherm model ‘endoR’ from NicheMapR (Kearney et al. 2021) to calculate the metabolic demands of common waxbills at each grid cell throughout Europe. This model estimates the metabolic heat production (QGEN) required for an animal to maintain a stable core body temperature, based on a heat balance approach. The model compares the simulated metabolic rate to a target rate and, if needed, allows for behavioural and physiological thermoregulation to achieve thermal balance. To account for increased metabolic heat production during active periods, we applied a daytime activity multiplier of 2.1 to the basal metabolic rate (BMR), consistent with empirical daytime metabolic rate (RMR) estimates for common waxbills (Sentís et al. 2023). Thus, the modelled metabolic heat generation was set to 2.1 × BMR during daylight hours and to BMR at night. If the simulated metabolic rate deviated by more than ±5% from the target range (an arbitrary error threshold accepted by the model), the model allowed animals to employ both physiological and behavioural thermoregulation (e.g. varying flesh thermal conductivity to simulate vasodilation or vasoconstriction, seeking shade) until a heat balance was achieved. The model thus calculated the necessary (in cold temperatures) metabolic rate that enabled an animal to maintain its body temperature within a tolerable range. Metabolic rates thereby represented hard limits for survival: if the simulated energy expenditure exceeded the estimated maximal energy intake, the animal was assumed to have died of hypothermia. The hourly outputs from the microclimate model (i.e. air temperature at reference and bird height, sky and surface temperatures, wind speed, relative humidity, and solar radiation) were used as inputs to the endotherm model.

Data on BMR and body mass were obtained from 237 wild common waxbills captured at two Portuguese sites in both summer and autumn, thus covering geographic and seasonal gradients within their invasive range (Sentís et al. 2025). Based on the average of these empirical measurements, BMR and body mass were set to 0.22 W and 8 g, respectively. Morphological traits that were difficult to measure on live individuals were taken from 340 museum specimens from the species’ native range, primarily from the Congo, but also Kenya, Tanzania, Cameroon, Uganda, Senegal, Rwanda, and South Africa (KBIN, Brussels; KMMA, Tervuren; MNHN, Paris). Feather characteristics were measured using a blunt probe accurate to 0.01 mm. For each specimen, feather depth (dorsal: 2.17 mm; ventral: 2.54 mm) and feather length (dorsal: 16.11 mm; ventral: 14.13 mm) were recorded on the torso. Measurements were taken twice at each location and averaged. For feather depth, the probe was inserted perpendicularly to the skin until it touched the surface; for feather length, it was aligned parallel to the feather orientation. The distance from the probe’s disappearance point to its tip was measured with a digital calliper.

Core temperature parameters were derived from body temperature measurements of 7 captive common waxbills, with the baseline core temperature set to 39.1 °C (reflecting body temperature at night, during rest) and the maximum allowable core temperature to 42.1 °C (reflecting maximum daytime body temperatures, Pacioni et al. 2023, Sentís et al. 2023). Birds’ body shape was modelled as an ellipsoid, and the maximum ratio of length to width was set to 3 (Sentís et al. 2023). Percentage of wet skin was set to 1% (Porter et al. 2006). Offset between air temperature and breath was set to 5 °C, and oxygen extraction efficiency was set to 25% (Kearney et al. 2021). Maximum panting rate and the multiplier on BMR at maximum panting level were set to 15 and 1, respectively (Kearney et al. 2021). All other NicheMapR parameter estimates were left at their default values.

### Model evaluation

#### Evaluating mechanistic models

When invasive range occurrences are available, forecasts of invasion risk are typically evaluated based on how well they predict locations that have already been invaded (Barbet-Massin et al. 2018). Correlative SDMs produce continuous habitat suitability values, ranging from 0 to 100, which reflect the probabilistic suitability of a given environment for the species. A wide range of evaluation metrics have been developed for assessing the performance of such models (Shabani et al. 2018). In contrast, mechanistic models do not yield probabilistic outputs. Instead, they predict metabolic costs based on environmental conditions. Mechanistic model outputs are therefore not expressed on a 0–100 suitability scale. Traditionally, these models are evaluated using ‘hard limits’: if the modelled metabolic requirements exceed a species’ allowable physiological thresholds, the location is deemed unsuitable and the species is assumed not to survive there. Here, we explore two complementary approaches for evaluating the forecast performance of our mechanistic model: (1) a traditional hard-threshold approach, and (2) a novel method that transforms the metabolic model outputs into a continuous 0–100 suitability scale, allowing for the application of standard SDM evaluation metrics.

For the hard-threshold method, if waxbills are unable to tolerate environmental conditions in even a single hour a year, that location could arguably be considered unsuitable. However, since NicheMapR incorporates a 5% model error (see above), we apply a more liberal threshold, designating a pixel as unsuitable only if more than 5% of the modelled hours indicate that environmental conditions exceed tolerance limits. The result is a binary suitability map, where each pixel is classified as either suitable or unsuitable. This map allows us to assess model sensitivity (i.e. the proportion of known invasive waxbill occurrences correctly predicted as suitable) and specificity (i.e. the model’s ability to correctly identify locations unsuitable for establishment; see below). Our second, continuous approach involves calculating the proportion of modelled hours per year in which metabolic costs fall within the waxbill’s physiological tolerance range. This yields values between 0 (no tolerable hours, i.e., conditions are unsuitable throughout the year) and 100 (all hours are tolerable), producing a continuous ’mechanistic suitability’ score to which traditional SDM evaluation metrics can be applied. This approach assumes that locations with a higher proportion of tolerable hours are more likely to support waxbill survival and establishment, and are therefore at greater risk of invasion.

To determine which environmental conditions are physiologically tolerable for waxbills, we defined a threshold based on the species’ metabolic expansibility (ME), which is based on the summit metabolism (Msum) (i.e. the maximum (rest) metabolic rate achieved during cold exposure (Cortés et al. 2015)), relative to its basal metabolic rate (BMR). In captive common waxbills, Pacioni et al. (2023) reported that Msum was, on average, 4.67 times the BMR. Accordingly, any hour in which the modelled metabolic cost exceeded this Msum threshold was considered unsuitable for waxbills.

#### Accessible areas for model evaluation

Evaluating invasive species distribution forecasts presents several challenges. One of the main issues is that while the presence of a species in the invasive range is generally a reliable indicator of environmental suitability - if a location were unsuitable, the species likely would not have established - the same cannot be said for locations where the species is absent. Absence may not reflect true unsuitability but rather the fact that the species has not yet had the opportunity to colonize those areas, for example due to dispersal limitations since its introduction. Given that all model evaluation metrics, in one way or another, rely on comparing presence with absence locations - or comparing areas used by the species to those available to it - model evaluation should ideally be restricted to the accessible area: the set of sites that have realistically been available to the species since its introduction, and that it could have colonized if conditions were suitable. Here, we explored two alternative definitions of the background area. First, we considered the whole of Europe as the invasive background, reflecting the assumption that pet bird trade has provided common waxbills with the opportunity to escape and establish populations across the continent. Second, we defined a more conservative accessible area based on known introduction locations and estimated invasion speeds. Using information from the literature, we identified sites where waxbills were successfully introduced in the Iberian Peninsula and calculated the Euclidean distance to the most distant confirmed waxbill records. Dividing this distance by the number of years between introduction and observation yielded an approximate colonization speed. For each successful introduction site, we then multiplied this speed by the number of years since introduction to generate a circular buffer representing the area that common waxbills could realistically have reached since their introduction. Because failed introductions may also provide valuable information about unsuitable areas, we incorporated data on such locations (from literature and transient GBIF occurrence records; see below). Around each site of known failed introduction, we applied an arbitrary 50 km buffer and then combined these with the dispersal-based buffers for the Iberian Peninsula to create a unified accessible area for Europe. All evaluation metrics were then calculated twice, one on the whole of Europe as accessible area, and once using a restricted ‘invasion history’ accessible area.

#### Invasive range occurrence data

We retrieved occurrence records for common waxbills from the Global Biodiversity Information Facility (GBIF) for all available records up to 2024. Only records with valid geographic coordinates were retained. To further refine the occurrences, we cross-checked occurrence points against the datasets of (Ascensão et al. 2021), the Global Avian Invasions Atlas (GAVIA) and the third Spanish Bird-Breeding Atlas III (SEO/BirdLife, 2022), which provide verified data up until 2020. To account for potential spread since 2020, we created a 80 km buffer around the presences documented by the atlases. Occurrences falling outside this buffer were considered more likely to be transient or incorrectly reported and were excluded. Finally, to limit sampling bias where multiple records were clustered in the same location, we aggregated the occurrence data at the 5 × 5 km resolution of our modelled suitability rasters. Each point was assigned to its corresponding raster cell, and any duplicates within the same cell were combined into a single occurrence.

#### Evaluation metrics

For our hard-threshold method (i.e. a pixel is considered unsuitable if more than 5% of the modelled hours indicate that environmental conditions exceed tolerance limits), model evaluation is measured using sensitivity and specificity. Model sensitivity ranges from 0 to 1 and reflects how many invasive waxbill occurrences are located in pixels predicted to be suitable by the model. Model specificity is the analogue for species absences, and in order to calculate it, a number of pseudo-absence locations equal to the number of invasive occurrences was drawn from the accessible area. Our model specificity values thus represent the number of pseudo-absence locations that our models indicate as unsuitable for the species. For our novel, continuous ’mechanistic suitability’ score, we applied three evaluation metrics commonly used for correlative SDMs. First, and similar to the hard-threshold method described above, we reclassified the 0 to 100 mechanistic suitability values into binary suitable/unsuitable categories to calculate sensitivity and specificity. However, unlike the hard-threshold approach, the threshold is not chosen a priori. Instead, we used the ecospat.max.tss function (Di Cola et al. 2017), which searches for the threshold that best balances sensitivity and specificity. It evaluates a range of possible thresholds and selects the one that results in an optimal trade-off between correctly predicting presences and absences. In addition, we calculated two so-called presence-only, threshold-independent metrics: the Boyce index and the partial AUC. The Boyce index assesses how much the model’s predictions differ from a random distribution of observed presences across the suitability gradient, with values ranging from −1 to +1. Positive values indicate that presences are concentrated in areas of high predicted suitability, values near zero indicate a random distribution, and negative values suggest predictions are worse than random (Hirzel et al. 2006). The partial AUC (area under the curve) is a modified version of the standard AUC that focuses on the most relevant part of the ROC curve—specifically the range where the model is expected to perform best. It places more emphasis on correctly predicting presences (i.e. higher sensitivity) while allowing some tolerance for false positives (lower specificity). Partial AUC values range from 0 (worse than random), through 1 (no better than random), to 2 (perfect discrimination) (Peterson et al. 2008, Cobos et al. 2019).

## 3. Results

When using the a priori hard-threshold (i.e. 5% model error, corresponding to a 95% cut-off in tolerable hours) to evaluate model predictive performance, the mechanistic model accurately identified invasive common waxbill occurrences across Iberia with a high sensitivity (0.985). Specificity was high, with a value of 0.854 when using the whole of Europe as accessible areas and moderate (0.470) when using an accessible area based on invasion history. Model evaluation based on metrics common to correlative distribution models similarly indicated a high model predictive performance. When iteratively searching for a cut-off value that balances sensitivity and specificity, we found an optimal cut-off value of 99% corresponding to a sensitivity of 0.938. Specificity ranged from 0.916 for the European accessible area definition to 0.686 for the invasion history area. When assessing the model using presence-only, threshold-independent evaluation metrics, the Boyce index showed a strong predictive discrimination across Europe (Boyce index = 0.989) but performed poorly within the accessible area based on invasion history (Boyce index = 0.052). Partial AUC ratios significantly exceeded random expectations in both background areas (Europe: partial AUC ratio = 1.829, p < 0.0001; accessible area: 1.412, p < 0.0001).

Southern Europe, particularly the Mediterranean region, was identified as having the highest invasion risk under current conditions (Figure 1). Under a +2 °C warming scenario, the model predicted that 28.4% of Europe would become climatically suitable (Figure 2; B), compared to 10.7% under current conditions (Figure 2; A). In this scenario, areas at risk of invasion expanded northwards into central Europe, with increased suitability projected in parts of France, Germany, and the Balkans. Under a +4 °C scenario, 67.7% of the area was classified as suitable, showing a marked expansion in climatic suitability across central and eastern Europe, including southern Scandinavia, while the northernmost latitudes and mountain ranges remained largely unsuitable (Figure 2; C).

**Figure 1.**
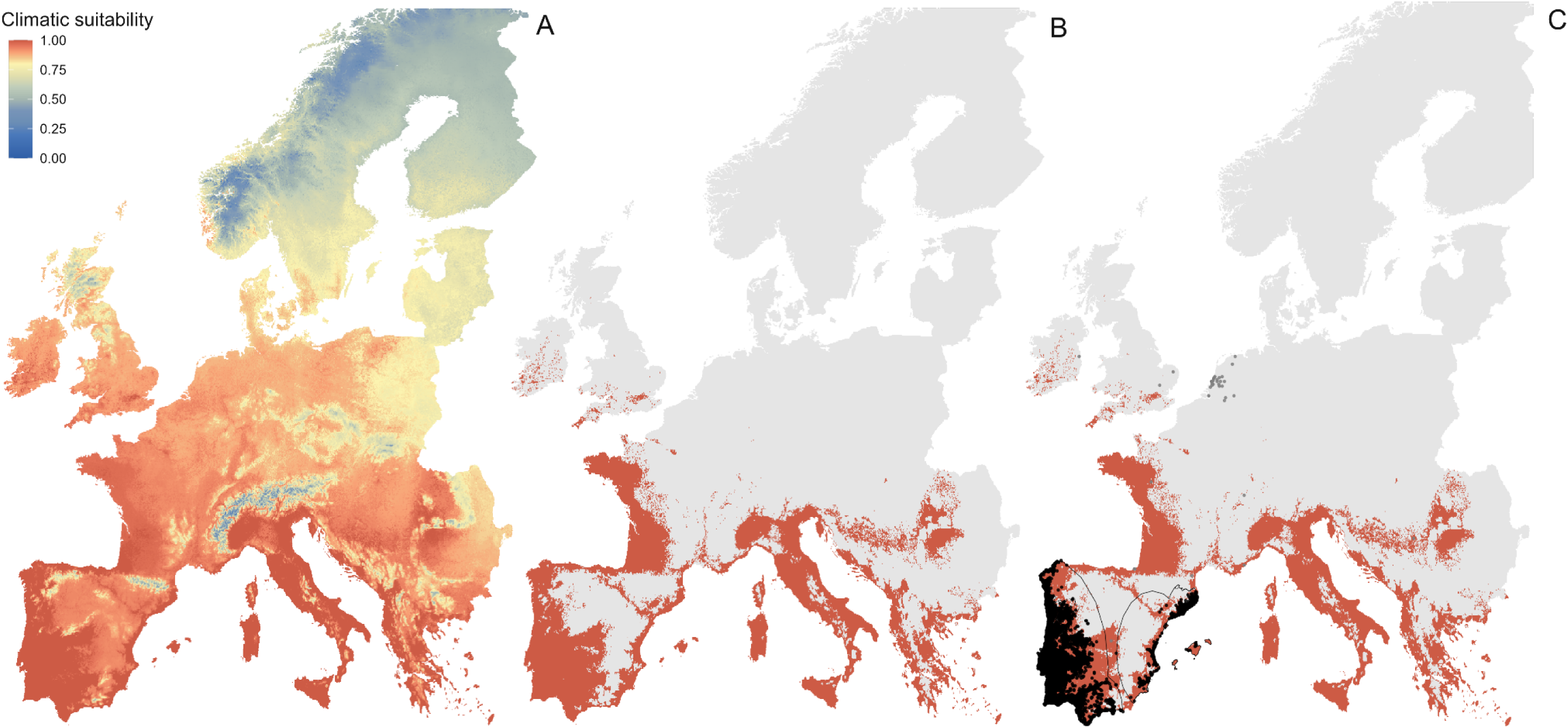
A. Predictions of climatic suitability for the common waxbill in Europe, values ranging from 0 (blue, low suitability) to 1 (red, high suitability). A value of 0 means that at that location, not a single hour per year is physiologically tolerable to common waxbills, a value of 1 means all hours are tolerable. B. Climatically suitable (red) versus unsuitable (grey) areas based on the 95% NicheMapR-derived cut-off. C. Same as panel B, with added occurrence records obtained through GBIF. Black dots indicate established populations and grey dots represent failed introductions. The estimated accessible area in Iberia based on invasion history is outlined in black.

**Figure 2.**
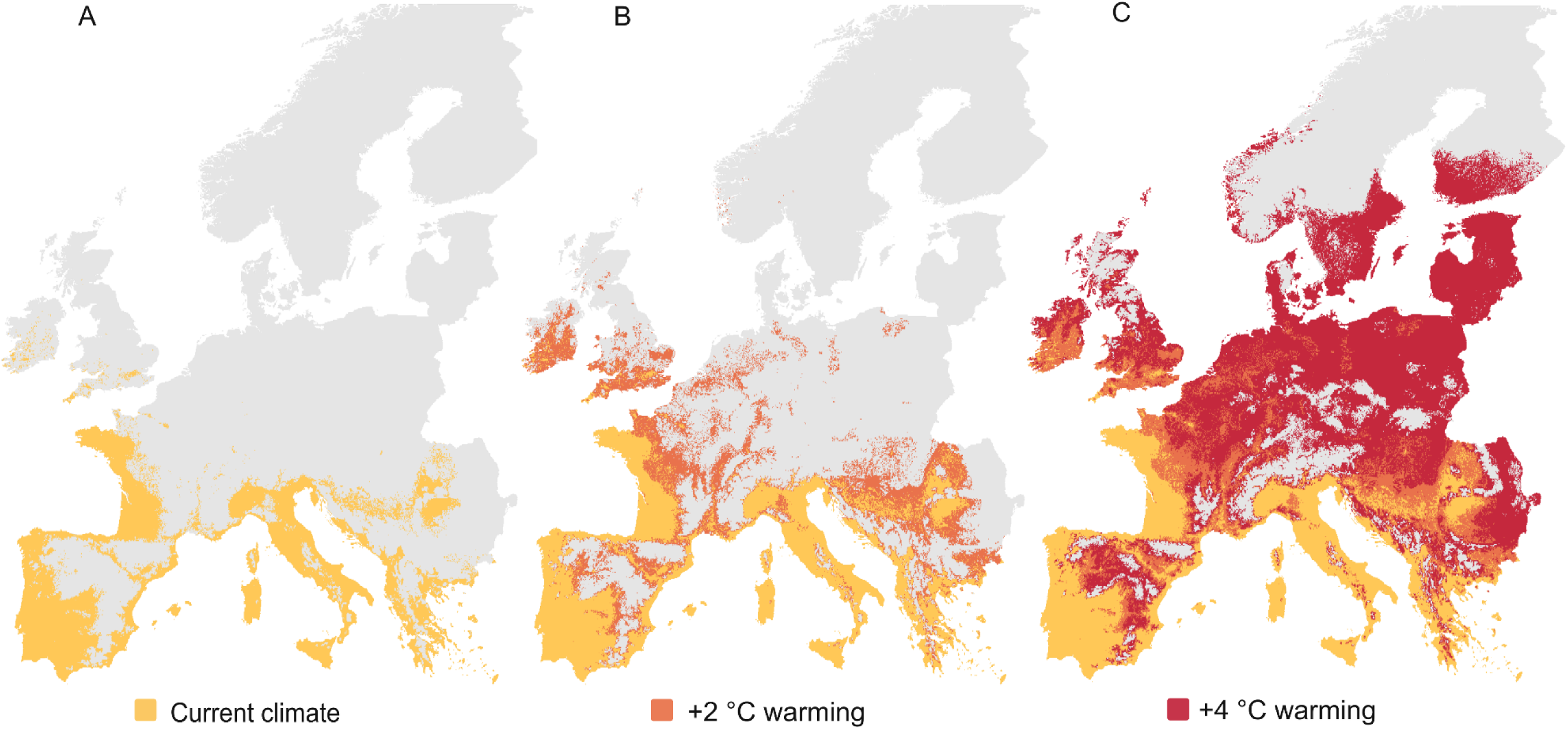
Projected climatic suitability of common waxbills in Europe. A. current climate; B. +2 °C warming scenario; C. +4 °C warming scenario. Yellow values indicate locations suitable under current conditions, orange indicates areas that become suitable under +2 °C warming and red the locations that become suitable under +4 °C.

## 4. Discussion

Preventing invasive species from establishing in new environments requires proactive strategies, yet predicting their potential geographic distribution remains a key challenge (Gallien et al. 2010, Araújo and Peterson 2012, Urban et al. 2016, Yates et al. 2018, Araújo et al. 2019, Battini et al. 2019). Correlative species distribution models (SDM) have become a cornerstone of invasion risk assessments due to their relatively low data requirements and ability to model distributions across large numbers of species using only occurrence records and environmental predictors (Guisan et al. 2014, Elith 2017, Srivastava et al. 2019). Indeed, their flexibility and scalability have made SDMs the method of choice in many large-scale ecological and conservation studies (Guisan et al. 2013, Engler et al. 2017, Nichols et al. 2024). However, when applied to invasive species, SDMs often face significant challenges (Liu et al. 2020, A. Lee-Yaw et al. 2022, Hui 2022, Young et al. 2025), highlighting the value of complementary approaches that are less dependent on occurrence data for model training. Here, we applied a biophysical model that integrates high-resolution physiological and microclimatic data to predict the current and potential future distribution of the common waxbill, one of Europe’s most successful invasive bird species. Our model successfully predicts the known distribution of the waxbill in the Iberian Peninsula and identifies additional regions in Europe that are currently at risk of invasion. Projections under a +2°C warming scenario suggest a moderate expansion, mainly into temperate western European areas and parts of the Balkans. A 4°C increase, however, predicts a much broader range expansion across Europe, including most of eastern Europe and southern Scandinavia.

### Strengths of a mechanistic, biophysical approach

Mechanistic models, such as the biophysical framework presented here, offer a promising alternative by directly linking species-specific physiological traits to environmental conditions (Kearney and Porter 2009), allowing spatially explicit predictions of where invaders can survive, even in climates where they have not yet been recorded. The common waxbill is instructive in this context, as it exemplifies many of the challenges and opportunities inherent in invasion modelling. As one of Europe’s most successful avian invaders, its spread has been the subject of numerous species distribution modelling efforts (see above). However, despite the relatively well-documented invasion history of this species, previous models have produced inconsistent results. Some provide reasonable predictions, while others fail to capture known aspects of the species’ range dynamics. These discrepancies are difficult to interpret, as they may be due to differences in the quality and resolution of occurrence data, the environmental predictors selected, or the specific modelling algorithms applied. Thus, the invasion of the common waxbill studied here highlights the potential for biophysical models to significantly improve the accuracy of invasion predictions and provide a more reliable basis for proactive invasive species management and broader conservation planning.

### Model performance and sources of uncertainty

While our model, by capturing the species’ basic climatic niche, achieves high sensitivity and accurately predicts most invasive presences, it may slightly overestimate habitat suitability, resulting in slightly lower specificity. In other words, it sometimes predicts suitable conditions in places where the species does not currently occur. This probably reflects the challenge of distinguishing between areas that the invasive species could have reached and those that it has reached. Although we attempted to account for this by evaluating model performance within an accessible area based on invasion history, this remains a crude approximation. Another reason is that the model only considers climatic variables. It therefore identifies climatically suitable areas without incorporating other ecological constraints, such as food availability, local habitats or biotic interactions, which can all prevent establishment even in otherwise suitable climates (Wisz et al. 2013, Peterson et al. 2015). Nevertheless, the model still shows moderate to good specificity, correctly rejecting many unsuitable areas, especially colder, central, and mountainous parts of Iberia. We also note that while the Boyce index is high when evaluated across the entire continent, it decreases when calculated only within the accessible area informed by invasion history. This is likely because presence-only metrics such as the Boyce index rely on contrasting suitability values between occupied and available areas. When the assessment area is restricted and lacks sufficient suitability variation, this contrast is diminished, reducing the discriminatory power of the metric (Hirzel et al. 2006).

A further consideration is the slight mismatches between occurrence data and predicted ranges. While sensitivity was high, a few known occurrences were not predicted as suitable. This small discrepancy may partly reflect limitations of the mechanistic model, as some of the traits used may not fully represent wild populations or capture intraspecific variation across environments. In addition, uncertainties in microclimate modelling and the use of relatively coarse spatial resolution may have masked microhabitat conditions enabling the species to occur in areas otherwise classified as unsuitable. On the other hand, not all occurrence records necessarily indicate established populations. Despite filtering, some GBIF records may represent vagrants, escapees, or short-term presences that do not reflect long-term population establishment.

### Thermal constraints on the current European range

Temperature appears to play a central role in shaping the invasion dynamics of the common waxbill in Europe. In Catalonia, for example, (Herrando et al. 2011) observed population fluctuations that coincided with the severity of winter conditions, suggesting sensitivity to low temperatures. Similarly, SDMs based on invasive range occurrences suggest that minimum winter temperatures limit the Iberian range of the species (Reino et al. 2009, Keller et al. 2020). These findings are consistent with recent evidence from Sentís et al. (2025), who showed that invasive waxbill populations in Iberia had reduced nutritional status and poorer feather quality compared to native (South African) birds, suggesting possible physiological stress near the species’ thermal limits. Indeed, Strubbe et al. (2023) hypothesized that established populations in Europe may occupy areas close to their upper thermoregulatory limits. While our biophysical model suggests that waxbills have colonized nearly all suitable habitats throughout Iberia, it also predicts potential for expansion along the Atlantic coast of western France and as far north as Brittany, likely due to the region’s relatively mild winters. Italy and other Mediterranean regions are identified as particularly vulnerable to future colonization (Figure 1). In these areas, waxbills are likely to benefit from a combination of mild winter temperatures, open habitats near water (Ribeiro et al. 2020), and the availability of peripheral niches in human-modified landscapes (Batalha et al. 2013). Their diet, consisting mainly of common grass seeds (Cardoso et al. 2018, Lucio-Puig et al. 2025), is readily available in many Mediterranean coastal habitats (Jiménez-Ruiz et al. 2021, Lázaro-Lobo et al. 2024). This convergence of favourable climatic conditions and likely abundant food resources creates a favourable ecological context for continued range expansion across most of southern Europe.

### Projected range expansion under climate warming

Under a warming climate, and assuming that water and food availability are not limited, our projections suggest that suitable areas for waxbills in Europe will expand. The modelled response is non-linear, with an increase of 2.7 times the current extent under a +2 °C scenario, and a larger expansion of approximately 6.3 times the current extent under a +4 °C scenario (Figure 2). This suggests that further warming could substantially reduce cold-related constraints and render currently marginal habitats suitable. This pattern is consistent with broader trends described by (Naimi et al. 2022), who suggest that as the climate warms and higher latitudes become ’tropicalised’, tropical birds may increasingly colonise temperate regions where cold winters have previously limited their establishment. Beyond lifting thermal constraints, climate change may also extend the duration of favourable seasons, increasing breeding opportunities and resource availability, thereby amplifying the potential for population growth and subsequent expansion (Halupka and Halupka 2017, Mingozzi et al. 2022, Halupka et al. 2023).

### Implications, caveats and future directions

Our biophysical model was parameterized using averaged species-specific measurements from wild-caught individuals and museum specimens, the latter providing morphological traits such as feather depth and length which are difficult to obtain from live birds, and it accurately predicted the species’ current invasive distribution. This suggests that morphological measurements derived from museum specimens can reliably provide estimates of key traits that cannot be collected in the field, and that models based on representative average trait values can generate informative, biologically meaningful predictions. Nevertheless, within-population variability likely exists and could further refine projections, particularly under scenarios involving novel climates or rapid evolutionary change (Kolbe et al. 2010, Pearman et al. 2010, Riddell et al. 2023, Fenollosa et al. 2025).

Additionally, the current framework does not incorporate upper thermal constraints, e.g. physiological limits beyond which performance declines due to heat stress and limited evaporative cooling capacity (Riddell et al. 2019). This could have led to an overestimation of climatic suitability under future warming scenarios, especially in the hottest Mediterranean regions where extreme heat events are expected to intensify (Carvalho et al. 2021). Under such extreme conditions, birds rely heavily on evaporative cooling, incurring substantial water losses to maintain thermal balance (Riddell et al. 2021). NicheMapR can simulate both cutaneous and respiratory evaporative water loss (Kearney et al. 2021). Comparing these outputs with critical thresholds of 11–15% body mass loss, beyond which dehydration becomes lethal in small birds (Wolf 2000, McKechnie et al. 2016), could be used to predict invasion risk in hot environments. Despite these limitations, our findings underscore that mechanistic SDMs grounded in high-quality, species-specific data are capable of providing robust forecasts of invasion dynamics and climatic suitability. This aligns with recent calls to expand the use of process-based models in invasion ecology (Strubbe et al. 2023, Fenollosa et al. 2025, Briscoe et al. 2023).

## Conclusions

Our findings demonstrate that mechanistic SDMs offer a powerful approach to forecasting invasion risk. For the common waxbill, the model captured both its current distribution and projected future expansion, identifying southern and western Europe as high-risk areas for invasion. As these regions are climatically suitable but currently unoccupied, early interventions such as trade restrictions and targeted surveillance may be critical to preventing future establishment, making them key targets for effective management. Overall, our results underscore the growing value of mechanistic models in invasion biology and their capacity to predict range expansions driven by climate change. More broadly, they highlight the potential for such models to be integrated into biodiversity forecasting frameworks to support proactive conservation planning and early detection of emerging invasion threats.

## Acknowledgments

We thank A Rocha Portugal, Preshnee Singh, and Ebrahim Ally for their support during fieldwork. We are also grateful to Marie Stessens, Ananya Agnihotri, and Lowie Tondelier for their assistance with museum data collection. Luís Reino was funded through the FCT contract ‘CEECIND/00445/2017’ under the ‘Stimulus of Scientific Employment - Individual Support’ and by FCT ‘UNRAVEL’ project 16000587 (PTDC/BIA-ECO/0207/2020). Marina Sentís acknowledges the support of FWO-Vlaanderen (project 11E1623N). This study also acknowledges funding by FWO-Vlaanderen (project G0E4320N) and by Methusalem Project 01M00221 (Ghent University) awarded to Frederick Verbruggen, Luc Lens, and An Martel.

